# Zmiz1 is required for mature β-cell function and mass expansion upon high fat feeding

**DOI:** 10.1101/2022.05.18.492530

**Authors:** Tamadher A. Alghamdi, Nicole A. J. Krentz, Nancy Smith, Aliya F. Spigelman, Varsha Rajesh, Alok Jha, Mourad Ferdaoussi, Jocelyn E Manning Fox, Han Sun, Zijie Sun, Anna L. Gloyn, Patrick E. MacDonald

## Abstract

Genome-wide association studies have identified hundreds of signals for type 2 diabetes (T2D), most of which confer risk through effects on gene expression. We previously identified the transcription factor *ZMIZ1* as a probable effector transcript in human islets, but how altered *ZMIZ1* expression impacts T2D risk is unknown. We now show that islets from carriers of the T2D-risk alleles have reduced islet insulin content and glucose-stimulated insulin secretion. To elucidate the mechanism for islet-cell dysfunction, we generated β-cell-specific *Zmiz1* knockout (Zmiz1^βKO^) mice. Male and female Zmiz1^βKO^ mice were glucose intolerant with impaired insulin secretion, compared with control littermates. Transcriptomic profiling of Zmiz1^βKO^ islets identified over 500 differentially expressed genes including those involved in β-cell function and maturity which we confirmed at the protein level. After high fat feeding, Zmiz1^βKO^ mice fail to expand β-cell mass and become severely diabetic. Thus, Zmiz1 is required for normal glucose homeostasis and may contribute to T2D risk by maintaining a mature β-cell state and allowing islet mass expansion upon metabolic stress.

## INTRODUCTION

Despite considerable progress in uncovering the genetic landscape for type 2 diabetes (T2D), progress in moving from a genetic association signal to effector transcripts and downstream biology remains slow (Krentz & Gloyn, 2020). Most genetic signals associated with T2D risk are located in non-coding regions, suggesting that they likely confer their risk through regulatory effects (Mahajan et al., 2022; Spracklen et al., 2020; Viñuela et al., 2020). We previously used expression quantitative trait locus (eQTL) analyses to identify T2D-risk variants which alter transcript expression in human islets and identified *ZMIZ1* as the likely effector transcript at a locus on chromosome 10 (van de Bunt et al., 2015).

ZMIZ1 (Zinc Finger MIZ-Type Containing 1) was first identified as a co-activator of the androgen receptor in human prostate epithelial cells (Sharma et al., 2003) and is a member of the Protein Inhibitor of Activated STAT(PIAS)-like family, a group of proteins that regulate transcription through several mechanisms including blocking DNA-binding of transcription factors, recruiting transcriptional coactivators or corepressors, and protein SUMOylation (Shuai & Liu, 2005). ZMIZ1 regulates several transcription factors including p53 (Lee et al., 2007), Smad3 (Li et al., 2006), and Notch1 (Pinnell et al., 2015). Complete deletion of *Zmiz1* in mice is embryonically lethal due to impaired vascular development (Beliakoff et al., 2008). How altered *ZMIZ1* expression contributes to T2D risk and influences insulin secretion remains poorly understood. We previously showed in human islets that overexpression of *ZMIZ1* reduced glucose-stimulated insulin-secretion whilst knockdown of *ZMIZ1* inhibited KCl-induced secretion (van de Bunt et al., 2015). Separately, knockdown of *ZMIZ1* in the human β-cell line EndoC-βH1 resulted in reduced insulin secretion and cell count (Thomsen et al., 2016).

In the current study we investigated the impact of T2D-associated variants at the *ZMIZ1* locus on β-cell function in islets from non-diabetic carriers, and characterised male and female mice bearing *Zmiz1* null β-cells. We demonstrate an association of T2D-risk alleles with reduced insulin content and glucose stimulated insulin secretion in human islets and a role for Zmiz1 in maintaining β-cell maturation, function, and expansion upon metabolic stress in mice.

## METHODS

### Human islet studies

Isolation of human islets and static glucose-stimulated insulin secretion assay have been described in the protocols.io repository (Lyon et al., 2019). DNA was extracted from exocrine tissue, spleen, or, if no other tissue was available, islets. Genotyping was performed on Illumina Omni2.5Exome-8 version 1.3 BeadChip array. Allelic imbalance was performed on ATAC-seq of 17 islets (Thurner et al., 2018) that were processed following the ENCODE ATAC-Seq pipeline (v1.9.3). Overlapping reads with the 10 SNPs in the 99% credible set (Mahajan et al., 2018) were remapped by WASP (v0.3.4) to correct mapping bias. SNPs and samples were further filtered based on the parameters (read depth >= 5 for each allele and heterozygous samples > 2) specified in (Thurner et al., 2018). Percentage of the reference allele counts were defined as REF counts / (REF counts + ALT counts) * 100. One sample t test was performed to test whether the ratio is different from the null allelic proportion of 0.5.

#### Animal studies

ROSA26 Cre reporter (also known as R26R) mice [B6;129S4-*Gt(ROSA)26Sor*^*tm1Sor*^/J] (Soriano, 1999) were obtained from the Jackson Laboratory (Bar Harbor, ME). Ins1-Cre knock-in mice (Thorens et al., 2015) were kept on a C57Bl/6J genetic background. *Zimz1*^*fl/fl*^ mice were generated by flanking coding exons 8 to 11 of *Zimz1* with two loxP sequences, and kept on C57Bl/6J genetic background. Ins1-Cre knock-in mice were crossed with *Zimz1*^*fl/fl*^ mice to generate the following cohorts of male and female mice: *Ins1*-*Cre*^*+*^*Zmiz1*^+/+^, *Ins1*-*Cre*^*+*^*Zmiz1*^*fl/+*^, and *Ins1*-*Cre*^*+*^*Zmiz1*^*fl/fl*^; referred to as Zmiz1^Ctrl^, Zmiz1^βHET^, and Zmiz1^βKO^, respectively. Mice were fed chow diet (5L0D*, PicoLab Laboratory Rodent Diet) until 12 weeks of age, after which time some mice were switched to high fat diet (HFD; 60 % fat; Bio-Serv, CA89067-471) for an additional 8 weeks.

Mice were fasted for 4-6 hours and had access to water prior to oral glucose tolerance test (OGTT) (1g/kg dextrose) (Smith, Ferdaoussi, et al., 2018) or intraperitoneal glucose tolerance test (IPGTT) (1g/kg dextrose) (Smith et al., 2019b) for chow fed groups (12 weeks of age). For HFD groups, 0.5g/kg dextrose was used for both OGTT and IPGTT (20 weeks of age). Blood was collected every 15 minutes for two hours and centrifuged at 4 °C for 10 minutes at 10,000 rpm. Glucose levels were measured using One Touch Ultra 2 glucometer (LifeScan Canada Ltd.; Burnaby, British Columbia, Canada), and plasma insulin levels were assessed using Insulin Rodent (Mouse/Rat) Chemiluminescence ELISA (cat# 80-INSMR-CH01, CH10, ALPCO, NH, USA). Insulin tolerance tests (ITTs) (Smith et al., 2019a) were performed by intraperitoneal injection of insulin 1 U/kg Humulin R (Eli Lilly) and blood glucose levels were assessed after the initial insulin delivery every 15 minutes for 2 hours.

For islet isolation and perifusion, pancreas of euthanized mice were perfused with collagenase(Smith, F Spigelman, et al., 2018). Isolated islets were hand-picked or subjected to purification using Histopaque Gradient Centrifugation (Smith, Lin, et al., 2018) and were cultured overnight. Glucose-stimulated insulin secretion was determined as previously described (Lin et al., 2021).

### β-Galactosidase Expression

Enzymatic X-gal staining of pancreas cryosections (5 μm thickness) was performed using an X-Gal Staining Kit (Cat#GX10003, Oz Biosciences INC, San Diego, CA, US) according to the manufacturer’s instructions. Briefly, cryosections were thawed and washed with 1X PBS. Sections were fixed with fixing buffer for 15 minutes at room temperature and then washed 2 times with 1X PBS. Freshly prepared 1X staining solution of X-Gal (5-bromo-4chloro-3-indoyl-*β*-D-galactopyranoside) was added to each section and incubated in a humidified environment at 37 °C overnight. The following day, slides were washed once with 1X PBS. ProLong Gold anti-fade reagent (P36930, Invitrogen) was applied, and slides were allowed to dry. Pancreatic sections were imaged under bright-field with an Olympus DP27 micrscope.

### Immunohistochemistry and β cell mass assessment

Mouse pancreas was weighed before fixing in Z-fix (VWR) and embedded with paraffin. Paraffin-embedded pancreatic tissue sections (3 μm thickness separated by 200 μm) were rehydrated and subjected to antigen retrieval with sodium citrate buffer (10mM Sodium citrate, 0.05% tween 20, pH 6.0) microwaved for 15 minutes at high temperature and allowed to cool for 20-40 minutes. 0.1% Triton-X (T9284, Sigma) was added for permeabilization for 5 minutes and slides were washed with 1X PBS 3 times for 5 minutes each. Tissues sections were then blocked with 20% goat serum (G9023, Sigma) for 30 minutes and stained with insulin antibody (dilution 1:5, IR002, Dako) for 1 hour at room temperature and washed 3 times with 1X PBS. Tissue sections were incubated with Alexa Fluor 488 goat anti-guinea pig (dilution 1:200, A11073, Invitrogen) secondary antibody for 1 hour at room temperature. Slides were then washed with 1X PBS 3 times and ProLong Gold anti-fade reagent with DAPI (P36931, Invitrogen) was applied. Slides were allowed to dry before imaging. Pancreatic sections were imaged at 10X objective using Zeiss COLIBRI Fluorescence Microscope and LED light source with 350-, 495-, or 555-nm filter set. For each animal, 3-4 sections were analyzed. Insulin-positive area was quantified with ImageJ software (NIH Image). The β cell mass was determined as the relative insulin-positive area of each section normalized to pancreas weight.

### RNA extraction, sequencing, and quantification

150 islets from chow-fed ZMIZ1^Ctrl^ and ZMIZ^β1KO^ mice were lysed in 1 mL of TRIzol Reagent (ThermoFisher Scientific) and RNA was extracted as per manufacturer’s guidelines. Library preparation and sequencing was performed by the Oxford Genomic Centre (Wellcome Centre for Human Genetics, Oxford, UK). All libraries were multiplexed and sequenced as 100-nucleotide paired-end reads. RNA libraries were sequenced to a mean read depth of 32 (±1.7) million reads per sample. STAR v2.5 (Dobin & Gingeras, 2015) was used to map reads to the mouse genome build GRCm38 with GENCODE m23 (ftp.ebi.ac.uk/pub/databases/gencode/Gencode_mouse/release_M23/gencode.vM23.annotation.gtf.gz) as the transcriptome reference. To quantify gene-level counts, featureCounts from the Subread package v1.5 was used (http://subread.sourceforge.net/) (Liao et al., 2014).

### PCA and differential expression analysis

Genes detected in all 14 samples at >1 count per million (cpm) were retained for downstream analysis, resulting in 12,047 protein-coding genes. Counts were normalised and transformed to log-cpm using the *voom* function within the *limma* package (v.3.32.5) (Ritchie et al., 2015) in R (v.3.3.2). Batch effect of islet isolations was corrected using *removeBatchEffect* in *limma* before principal component analysis. DESeq2 (v1.26.0) (Love et al., 2014) was used to identify 556 differentially-expressed genes (padj <0.05) between control and knockout islets. For sex-specific differences, the above analysis was performed on male or female only samples. The online tool Integrated System for Motif Activity Response Analysis (ISMARA) was used for computationally predicted regulatory sites for transcription factors (https://ismara.unibas.ch/mara/) (Balwierz et al., 2014).

### Western Blotting

Isolated islets were cultured overnight in RPMI with 11.1 mM glucose. Islets (n=100-200) were picked washed once with 1X Dulbecco’s Phosphate Buffered Saline (14190-144; gibco) PBS and collected in 30 μl of cell lysis buffer (C2978, Sigma) supplemented with protease inhibitor cocktail (P8340, Sigma). Islets were sonicated by using Heat Systems-Ultrasonics W-385 Sonicator Ultrasonic Processor at 4 °C. Protein concentration was estimated by Quick Start Bradford 1 X Dye reagent (5000205, BioRad) using a microplate reader (EnVision Multilabel Plate Reader, PerkinElmer). A non-reducing lane marker sample buffer (39001, Thermo Scientific) was added to a total of 10 μg of protein from islet cell lysates and the mixture was heated for 5 minutes at 95 °C. Samples were then loaded, separated by SDS-PAGE (10% gel), and transferred to PVDF Immobilon-P transfer membrane (IPVH00010, Millipore). Membranes were blocked by 5% milk in Tris-buffered saline with 0.1% tween 20 (TBST) for 1 hour at room temperature, then probed with primary antibodies in 5% bovine serum albumin (BSA) in TBST overnight at 4°C with the following concentrations: Zmiz1(1:1000, Santa Cruz sc-367825), CD81 (dilution 1:1000; Cell Signaling 10037S), Aldh1a3 (dilution 1:1000, Novus biologicals NBP2-15339), β-actin (dilution 1:2000; Santa Cruz sc-47778). The following day, membranes were washed three times with 1X TBST and incubated with secondary antibodies in 5% milk in TBST for one hour at room temperature with the following concentration of secondary antibodies: donkey anti-rabbit IgG (1:5000, NA934, GE Healthcare) and goat anti-mouse IgG (1:5000, 115-035-146, Jackson ImmunoResearch). Membranes were washed three times with 1X TBST and incubated with ECL (45002401, GE Healthcare) for 5 minutes. Protein bands were imaged by ChemiDoc imaging system (Bio-Rad).

### Statistical analysis

Data are expressed as means±SEMs. Statistical significance was determined by one-way ANOVA followed by Tukey’s multiple comparisons test or two-way ANOVA followed by Bonferroni post-test to compare means between groups. Unpaired t test was used for comparison between two groups. Statistical analyses were performed using GraphPad Prism 9 for Mac OS X (GraphPad Software Inc., San Diego, CA).

### Approvals

Donor organs from individuals without type 2 diabetes were obtained with written consent and approval of the Human Research Ethics Board of the University of Alberta (Pro00013094; Pro 00001754). All animal studies were approved by the Animal Care and Use Committee at the University of Alberta (AUP00000291, AUP00000405).

## RESULTS

### T2D risk alleles of *ZMIZ1* rs703972 and rs12571751 are associated with reduced insulin content and secretion in primary human islets

We assessed secretory phenotypes in isolated islets from carriers of T2D risk variants at *ZMIZ1* in a cohort of 232 donors without diabetes (**Figure 1**). There are two independent signals at the *ZMIZ1* locus, one of which has been fine mapped to a ∼11kb interval and contains 10 SNPs in the credible set including the lead SNP rs703972 and the previously reported eQTL rs12571751 **(Figure 1A)**. The sex distribution, age, body mass index, and HbA1c were not different among genotypes (**Suppl Figure 1**). Islets from homozygous carriers of the T2D-risk alleles at rs703972 (G allele) and rs12571751 (A allele) had significantly lower insulin content compared to noncarriers (**Figure 1B-C**). Although no differences were observed in insulin secretion at low glucose (1 mM) concentrations, there was a reduction in insulin secretion in response to high glucose (16.7 mM), in the T2D risk-allele carriers (**Figure 1D-E**). The second signal has been fine mapped to an ∼834kb interval containing 861 variants in the credible set. There was no similar effect of the lead SNP (rs1317617) at this second locus, which may point to another effector gene at this signal (**Suppl Fig 1**).

**Figure 1.**
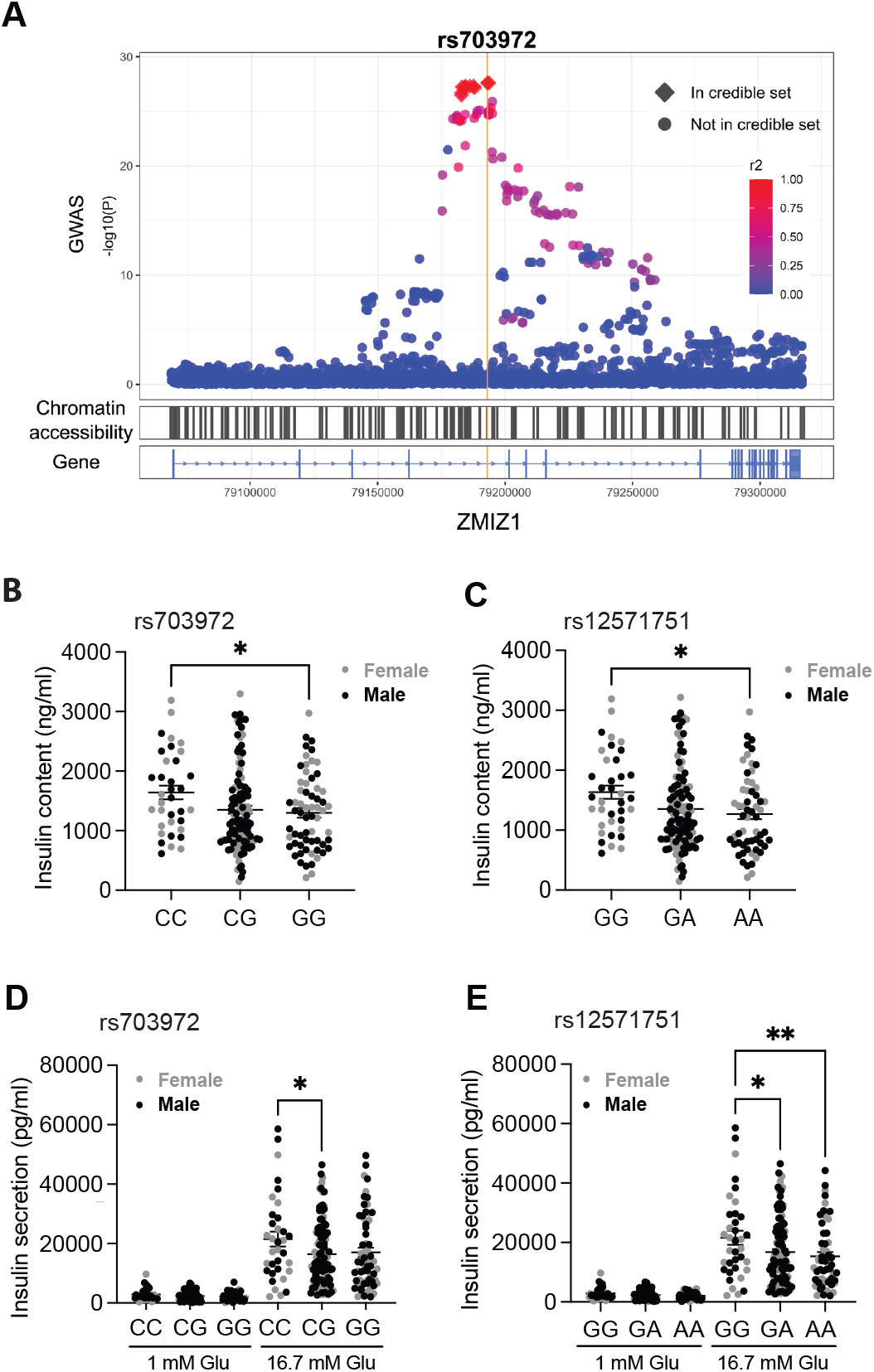
Effect of the T2D-associated alleles in *ZMIZ1* on insulin secretion in primary human islets. **(A)** Locus zoom plot for the *ZMIZ1* locus. The 10 SNPs in the 99% credible set defined by Mahajan et al (Mahajan et al., 2018) are shown as diamonds. LD r2 values are from the European population in TopLD. The lead SNP (rs703972) at the signal is shown with the yellow line. Chromatin accessibility in human islets is shown with open chromatin indicated by black lines. **(B-C)** Total insulin content in human islets from genotyped human donors. **(D-E)** Insulin secretion in response to low glucose (1 mM) or high glucose (16.7 mM) by *ZMIZ1* genotype at the lead SNP (rs703972) and previously reported eQTL (rs12571751). rs703972 CC, n=37; CG, n=126; GG, n=69 where G is the risk allele. rs12571751 GG, n=39; GA, n=127; AA, n=66 where A is the risk allele. Sex, age, BMI and HbA1c distribution is shown in Supplementary Figure 1. Glu, glucose. Data are mean ± SEM. ^**\***^*P*<0.05, ^**\*\***^*P*<0.01 by one-way ANOVA followed by Tukey’s multiple comparisons test.

To understand how T2D risk alleles at the first signal could alter *ZMIZ1* expression and to determine which SNP is functional at the locus we explored allelic differences in chromatin accessibility for credible set variants. We accessed publically available ATAC-seq data from 17 human islet donors and identified heterozygous individuals for three (rs703977, rs12571751, and rs810517) credible set variants which met our criteria (read depth >= 5 for each allele and >2 heterozygous donors) for inclusion in the analysis. No heterozygous carriers of the lead SNP rs703972 had sufficient read depth for inclusion in the analysis. After correcting for potential mapping biases using WASP (van de Geijn et al., 2015), there was a significant imbalance in allelic-specific chromatin accessibility for rs703977 (p=0.02) with the T2D-risk allele (T) having more open chromatin (**Table 1**), which would suggest increased *ZMIZ1* expression at the locus and is consistent with our previously published eQTL analysis (van de Bunt et al., 2015).

**Table 1:** Allelic imbalance in open chromatin of T2D-associated variants in *ZMIZ1*.

| Variant | DIAGRAM<br>P-value | Fgwas<br>T2D<br>PPA | Allelic<br>imbalance<br>Allele<br>Ratio<br>(Allele #) | Allelic<br>imbalance<br>WASP<br>P-value | T2D<br>risk<br>allele | Chromatin<br>state of<br>risk allele |
| --- | --- | --- | --- | --- | --- | --- |
| rs12571751 | 3.3E-27 | 1.5E-02 | 0.43<br>(52 A VS<br>68 G<br>alleles) | 0.59 | A | No<br>difference |
| rs703977 | 5.0E-28 | 1.1E-01 | 0.65 (95 T<br>VS 52 G<br>alleles) | 0.02 | T | ↑ Open |
| rs810517 | 2.0E-27 | 2.6E-02 | 0.49 (49 C<br>VS 52 T<br>alleles) | 0.65 | C | No<br>difference |

### Deletion of *Zmiz1* from pancreatic β-cells results in impaired glucose tolerance in mice

To characterize the role of pancreatic β-cell *Zmiz1* on glucose homeostasis, we selectively deleted the *Zmiz1* in β-cells by crossing Ins1-*cre*^*+*^ (Thorens et al., 2015) with *Zimz1*^fl/fl^ mice (**Figure 2A**). *Zmiz1*^*fl/fl/*^*:Ins1-Cre*^+^, *Zmiz1*^*fl/+*^*:Ins1-Cre*^+,^ and *Zmiz1*^*fl/fl/*^*:Ins1-Cre*^+^ littermates were generated, and are referred to as Zmiz1^Ctrl^, Zmiz1^βHET^, and Zmiz1^βKO^, respectively. All mice, including Zmiz1^Ctrl^, had Cre recombinase expression in β-cells and was confirmed by breeding the Ins1-*cre*^+^ knock-in mice with ROSA26 Cre reporter mice (*R26R*^fl/fl^) (Soriano, 1999) followed by X-gal staining (**Suppl Figure 2A**). Loss of Zmiz1 in Zmiz1^βKO^ islets was confirmed by immunoblotting (**Figure 2B**). We assessed OGTT, IPGTT and ITT in 12 week old mice on a chow diet (**Figure 2C**). Both female and male Zmiz1^βKO^ mice were glucose intolerant compared with Zmiz1^Ctrl^ littermates (**Figure 2C-F**). Compared with littermate controls, insulin tolerance of female and male mice was not different across all groups (**Suppl Figure 2B-C**), suggesting β-cell specific loss of Zmiz1 does not affect insulin sensitivity. Glucose-stimulated insulin secretion from isolated islets was markedly reduced from both the female and male Zmiz1^βKO^ mice (**Figure 2G-H**).

**Figure 2:**
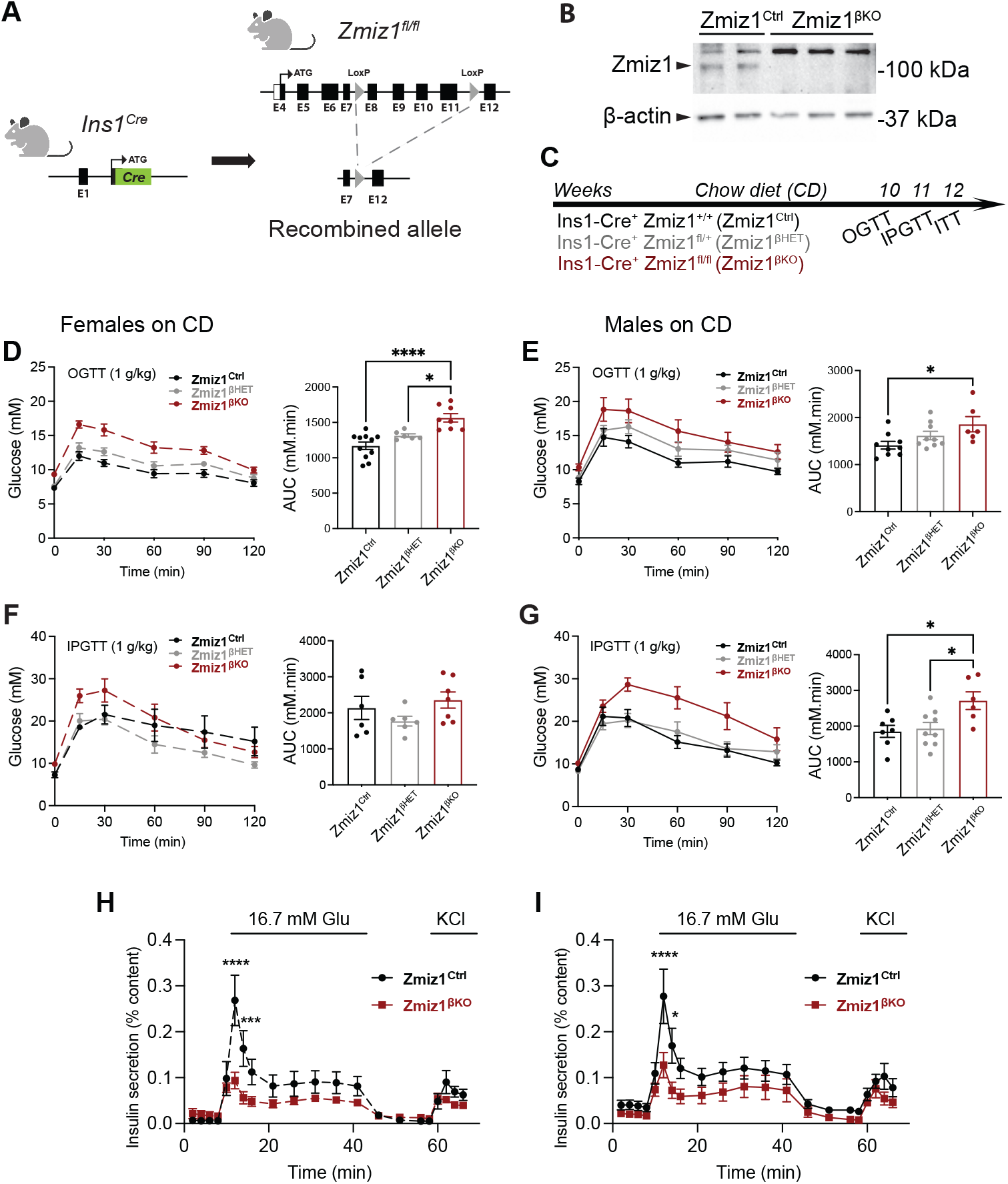
Loss of ZMIZ1 in β-cells impairs glucose homeostasis. **(A)** Schematic of *Ins1*^Cre^ knock-in allele, Zmiz1 floxed allele, and the recombined allele. **(B)** Immunoblotting for Zmiz1 (100 kDa) and β-actin (43 kDa) in lysates of primary islets isolated from Zmiz1^Ctrl^ and Zmiz1^βKO^ mice. **(C)** Schematic of experimental workflow on chow diet fed mice. **(D-E)** Oral glucose tolerance test (OGTT) and **(F-G)** intraperitoneal glucose tolerance test (IPGTT) in female (n=6-11 per group) and male (n=7-9 per group) Zmiz1 knockout and control mice. **(H-I)** Insulin secretion from female **(H)** and male **(I)** Zmiz1^Ctrl^ or Zmiz1^βKO^ mouse islets in response to glucose (n= 6-10 per group). CD, chow diet. Glu, glucose. AUC, area under the curve. Data are mean ± SEM. ^**\***^*P*<0.05, ^**\*\*\*\***^*P*<0.0001 by one-way ANOVA followed by Tukey’s multiple comparisons test or by two-way ANOVA followed by Bonferroni post-test.

### *Zmiz1* loss causes dysregulation of genes implicated in β-cell function and maturation

RNA-sequencing was performed on islets isolated from male and female Zmiz1^βKO^ and Zmiz1^Ctrl^ mice at 12 weeks of age (**Figure 3A**). There were 295 upregulated and 265 downregulated genes in Zmiz1^βKO^ islets (padj <0.05) (**Figure 3B**). Key β-cell maturity markers were downregulated (*Mafa, Glp1r, Slc2a2, Nkx6-1, Ins2, Ins1*) (Salinno et al., 2019), as were genes with known roles in insulin secretion and glucose metabolism (**Figure 3C**). Several genes implicated in β-cell proliferation and differentiation were also dysregulated (*E2f3, Rfx3, Nfatc1, E2f1*) (**Figure 3C**) (Ait-Lounis et al., 2010; Fajas et al., 2004; Rady et al., 2013; Simonett et al., 2021). Intriguingly, markers of immature or dedifferentiated β-cells, including *CD81* (Salinno et al., 2021) and *Aldh1a3* (Kim-Muller et al., 2016), were upregulated in the Zmiz1^βKO^ islet transcriptome (**Figure 3C**). Immunoblotting confirmed upregulation of CD81 and Aldh1a3 protein in Zmiz1^βKO^ islets (**Figure 3D-E**), highlighting a potential role for Zmiz1 in maintaining β-cell maturation.

**Figure 3.**
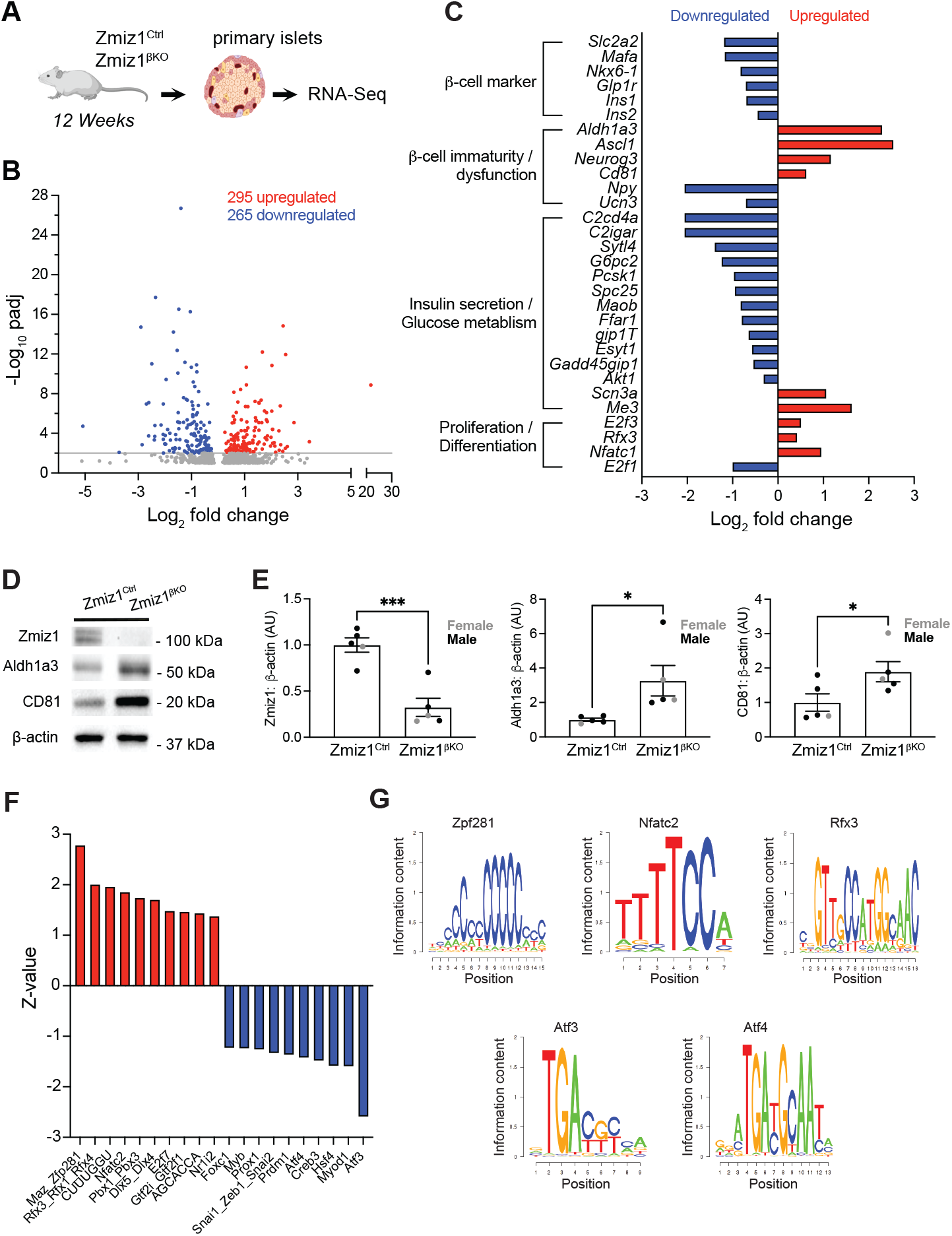
*Zmiz1* loss in β-cells results in dysregulation of genes implicated in β-cell function and maturation. **(A)** Schematic of experimental workflow of RNA-seq of mouse islets from female Zmiz1^Ctrl^ (n=3), female Zmiz1^βKO^ (n=4), male Zmiz1^Ctrl^ (n=4), and male Zmiz1^βKO^ (n=3) mice. **(B)** Volcano plot of the differentially expressed genes in Zmiz1^βKO^ islets. **(C)** Selected differentially expressed genes from the RNA-seq data and their known cellular functions. **(D)** Immunoblotting for Zmiz1, Aldh1a3, CD81, and β-actin in lysates of primary islets from Zmiz1^Ctrl^ or Zmiz1^βKO^ mice. **(E)** Quantification of band intensity of the indicated proteins in lysates of primary islets from Zmiz1^Ctrl^ (n=5) or Zmiz1^βKO^ (n=5) mice. **(F)** ISMARA analysis of top transcription factor binding motifs based on their motif activity (Z-value). **(G)** Selected predicted transcription factor binding motifs displayed. Data are mean ± SEM. ^**\***^*P*<0.05, ^**\*\*\***^*P*<0.001 by student t test. Upregulated (red). Downregulated (blue).

To further understand the transcriptional network which Zmiz1 regulates in β-cells, we performed *in silico* analysis to identify the potential transcription factors downstream of Zmiz1. Integrated Motif Activity Response Analysis (ISMARA) uses RNA-seq data to predict key transcription factors driving the observed changes in gene expression (**Figure 3F**). Among the top-ranked predicted regulatory motifs sorted by activity significance (Z-value) are transcription factors implicated in β-cell proliferation (*Nfatc2*), cell differentiation (*Zfp281* and *Rfx3*), and β-cell function and maturation (*Atf3* and *Atf4*) (**Figure 3F-G**).

### Loss of *Zmiz1* restricts β-cell expansion upon high fat feeding, leading to severe glucose intolerance

We next examined the effect of *Zmiz1* deletion from β cells upon high-fat feeding. 12-week old male and female Zmiz1^βKO^ mice and controls were put on high fat diet (HFD) for 8 weeks and OGTT, IPGTT, and ITT were performed (**Figure 4A**). Both female and male HFD-Zmiz1^βKO^ mice developed fasting hyperglycemia and severe glucose intolerance by 20 weeks of age (**Figure 4B-C**). In males, glucose intolerance measured by IPGTT (**Figure 4D**) was more obvious than female HFD-Zmiz1^βKO^ mice (**Suppl Figure 3A**) where we observed much less induction of insulin intolerance (**Suppl Figure 3B**) than in male mice (**Figure 4E**).

**Figure 4.**
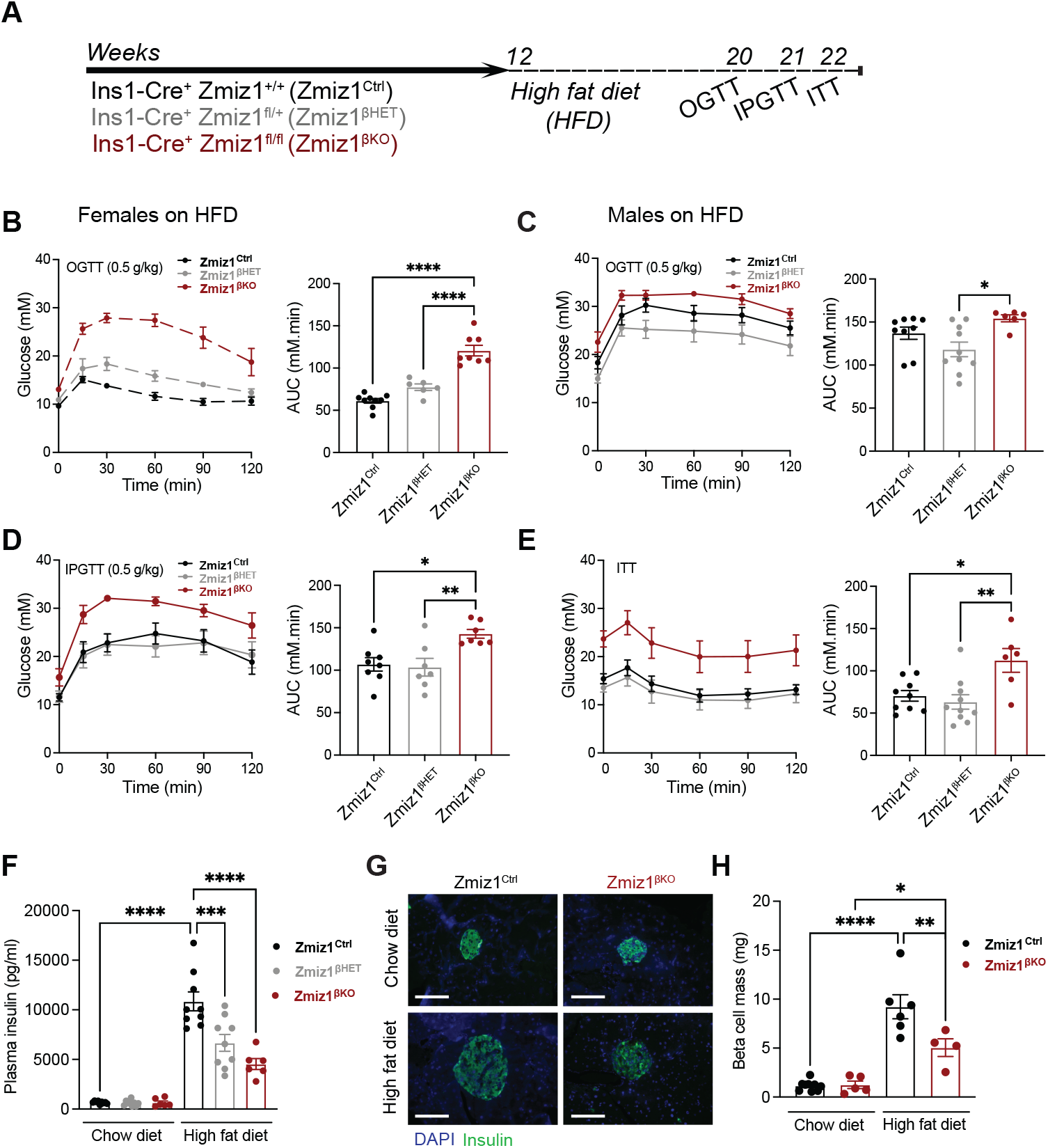
Loss of *Zmiz1* in β-cells reduces β-cell mass and worsens impaired glucose tolerance in mice challenged with high fat diet. (**A)** Schematic of experimental workflow on high fat diet (HFD) fed mice. **(B)** Oral glucose tolerance test (OGTT) in female Zmiz1 knockout and control mice (n=6-9 per group). **(C-F)** Oral glucose tolerance test (OGTT) **(C)**, intraperitoneal glucose tolerance test (IPGTT) **(D)**, insulin tolerance test (ITT) **(E)**, plasma insulin following OGTT **(F)**, in male control and Zmiz1 knockout mice (n=6-10 per group). **(G)** Representative immunostaining image of β-cell mass in Zmiz1^Ctrl^ and Zmiz1^βKO^ mice fed with HFD (scale bars= 100 μm). **(H)** β-cell mass relative to pancreas weight in Zmiz1^Ctrl^ (n=6) and Zmiz1^βKO^ (n=4) mice following HFD. Average of 3-4 pancreatic sections per animal was considered one replicate. Insulin (green), nuclei (blue). HFD, high fat diet. AUC, area under the curve. Data are mean ± SEM. ^**\***^*P*<0.05, ^**\*\***^*P*<0.01, ^**\*\*\***^*P*<0.001, ^**\*\*\*\***^*P*<0.0001 by one-way ANOVA followed by Tukey’s multiple comparisons test.

Fasting insulin is increased upon HFD in both males (**Figure 4F**) and females (**Suppl Figure 3C**), and this was blunted in the male HFD-Zmiz1^HET^ and HFD-Zmiz1^βKO^ mice (**Figure 4F**). To investigate the underlying mechanism, we assessed β-cell mass in HFD-Zmiz1^βKO^ mice and littermate controls. The female HFD-Zmiz1^Ctrl^ mice did not show an expansion of β-cell mass following the 8-week HFD (**Suppl Figure 3D-E**), consistent with the resistance of female mice to HFD-induced glucose intolerance (Pettersson et al., 2012) and insulin resistance (**Suppl Figure 3B**). As such, we see little difference in β-cell mass in the female HFD-Zmiz1^βKO^ mice (**Suppl Figure 3D-E**). In males however, while β-cell mass is not different upon chow feeding, the expansion seen with HFD in controls is blunted in male HFD-Zmiz1^βKO^ mice (**Figure 4G-H**). Glucose-stimulated insulin secretion was not different in both male and female HFD-Zmiz1^βKO^ mice compared to control (**Suppl Figure 3F-G**). Taken together, these observations suggest a role for Zmiz1 in β-cell mass expansion upon metabolic stress.

## DISCUSSION

Recent efforts to identify causal variants associated with T2D risk require parallel efforts to identify the mechanisms underpinning these association signals that will be crucial for accelerating translation of T2D genetic discoveries into clinical applications. In the current study, we characterized the functional role of the T2D risk gene *ZMIZ1* in pancreatic β-cells.

In humans, we demonstrate that islets from carriers of T2D risk alleles at the *ZMIZ1* locus have reduced insulin content and insulin secretion in response to high glucose compared with noncarriers, suggesting a key role for ZMIZ1 in β-cell function. Efforts to provide a direction of effect at the *ZMIZ1* locus through islet genomics continue to support a role for increased expression as a cause for elevated diabetes risk: allelic imbalance of chromatin accessibility at rs703977 supports the T2D-risk allele impacting islet function through increased *ZMIZ1* expression. The identification of chromatin QTLs at this locus in larger numbers of islet samples would strengthen this observation. Our findings are in line with our previous studies demonstrating that perturbation of *ZMIZ1* expression in human islets and β-cells negatively influences insulin secretion (Thomsen et al., 2016; van de Bunt et al., 2015). In human islets, both *ZMIZ1* overexpression and knockdown reduced insulin secretion (van de Bunt et al., 2015), although the effect of partial knockdown (40% of control) was modest, resulting in a ∼10% reduction in insulin secretion in response to KCl-stimulation. In a follow up study, knockdown of *ZMIZ1* in the human β-cell line EndoC-βH1 lead to a reproducible reduction in insulin secretion and cell count (Thomsen et al., 2016). Taken together, these data suggest that *ZMIZ1* levels are precisely regulated to support robust insulin secretion. The relatively modest *ZMIZ1* knockdown in human islets (van de Bunt et al., 2015) could account for the differences observed between those experiments and the Zmiz1^βKO^ mice. Alternatively, the acute knockdown of *ZMIZ1* may have been of insufficient duration and at an inappropriate time point to observe a clear secretory defect, given current data supporting a role for Zmiz1 in β-cell maturation.

In chow-fed mice, β-cell specific deletion of *Zmiz1* results in glucose intolerance with no change in insulin sensitivity, highlighting an important role of Zmiz1 in maintaining glucose homeostasis. In chow-fed animals, impaired glucose tolerance following loss of β-cell *Zmiz1* likely results from reduced glucose-stimulated insulin secretion as β-cell mass was unaffected in the Zmiz1^βKO^ mice. Our RNA-seq analysis showed that *Zmiz1* absence in β-cells results in downregulation of mature β-cell markers and upregulation of immature and dedifferentiated β-cell markers including *Aldh1a3* and *Cd81* (**Figure 3C**). The enzyme aldehyde dehydrogenase 1 isoform A3 (ALDH1A3) has been shown to mark dysfunctional β-cells that have progenitor like features including the expression of *Neurgo3* (Kim-Muller et al., 2016), which itself was upregulated in Zmiz1^βKO^ islets. CD81 has been recently identified as a novel surface marker for immature and de-differentiated β-cells in the adult mouse and human islets (Salinno et al., 2021, p. 81). Salinno et al. showed that subpopulation of high CD81 expressing mouse β-cells have low expression levels of the mature β-cell marker *Ucn3*, which we also find to be downregulated in the Zmiz1^βKO^ islets. Moreover, a similar expression pattern of *Cd81* and *Aldh1a3* was reported in STZ-diabetic mouse β-cells (Salinno et al., 2021). In our study, we further confirmed the upregulation of both CD81 and ALDH1A3 at the protein level in adult Zmiz1^βKO^ islets, which demonstrates a role for Zmiz1 in β-cell maturation.

The implication of Zmiz1 in β-cell maturation is further supported by the upregulation of the transcription factor *Rfx3* in Zmiz1^βKO^ islet transcriptome profile and in the ISMARA analysis. This is likely a compensatory effect since RFX3 plays an essential role in the differentiation and function of mature β-cells (Ait-Lounis et al., 2010). Although it is not clear how Zmiz1 contributes to the establishment or maintenance of β-cell maturity, our ISMARA analysis implicates the transcription factors ATF3 and ATF4, master regulators of the cellular stress response with dual roles in glucose homeostasis (Juliana et al., 2018; Ku & Cheng, 2020, p. 3; Wortel et al., 2017). The role of ATF3 is tissue-specific and context-dependant (Ku & Cheng, 2020). Several studies reported a protective role of ATF3 in β-cells (Gurzov et al., 2012; Zmuda et al., 2010). ATF3 deficiency in HFD-fed mice exacerbates glucose intolerance and impairs insulin secretion without affecting β-cell mass (Zmuda et al., 2010). ATF3 is known to be regulated by ATF4 (Wortel et al., 2017), which has recently been shown to play a protective role in diabetic Akita mice by preserving β-cell identity (Kitakaze et al., 2021). β-cell specific deletion of *Atf4* in Akita mice resulted in β-cell dedifferentiation (Kitakaze et al., 2021). However, these observations were hardly seen in β-cell specific *Atf4* knockout mice although β-cell proliferation was markedly reduced in these mice (Kitakaze et al., 2021).

A role of Zmiz1 in T cell-development by regulating Notch signalling has been previously reported (Pinnell et al., 2015). Although Notch signalling plays a role in β-cell maturation (Bartolome et al., 2019; Dror et al., 2007), altered expression of Notch target genes were not observed in the Zmiz1^βKO^ islet transcriptome. In zebrafish, inactivating CRISPR/Cas9 mutations in the zmiz1a gene in zebrafish results in lethality at 15 days post fertilization and delayed erythroid maturation, demonstrating a crucial role of Zmiz1a in terminal differentiation of erythrocytes (Castillo-Castellanos et al., 2021). The authors also showed that loss of Zmiz1a in zebrafish caused a dysregulation in autophagy (Castillo-Castellanos et al., 2021). Dysregulation of autophagy genes was not observed in Zmiz1^βKO^ islet transcriptome with the exception of the autophagy gene *Atg4d*, which was downregulated. Thus, although Zmiz1 has been implicated in cellular development through Notch and autophagy pathways, we find no evidence supporting these pathways within the islet.

Among the potential Zmiz1 binding transcription factors predicted by ISMARA analysis is the suppressor Znf281, a zinc-finger transcription factor that regulates embryonic stem cells differentiation by acting as a repressor of many stem cell pluripotency genes (Fidalgo et al., 2011). Znf281 has been shown to inhibit differentiation of cortical neurons (Pieraccioli et al., 2018). Accordingly, a possible mechanism by which Zmiz1 maintains β-cell maturity is through activation of transcriptional suppressors of genes associated with de-differentiated cell state such as Znf281.

Severe glucose intolerance was observed, particularly in response to oral glucose, in both male and female Zmiz1^βKO^ mice subjected to HFD-feeding for 8 weeks. The worsened glucose intolerance in response to oral compared to IP glucose suggests an implication of incretin response, which is known to be enhanced following HFD (Ahrén et al., 2008; Gupta et al., 2017; Yamane et al., 2016). This is consistent with our previous observations in HFD-mouse models (Lin et al., 2021).

In male Zmiz1^βKO^ mice, HFD feeding resulted in impaired β-cell mass expansion and reduced fasting insulin levels compared to controls. Notably, Nfatc1/2, among the differentially expressed genes and the top-ranked Zmiz1-binding candidate, is known to promote β-cell proliferation in mouse and human islets (Keller et al., 2016; Simonett et al., 2021). The β-cell mass phenotype was less obvious in females, and accordingly the fasting hyperglycaemia was much more pronounced in males. This is not necessarily due to sex-differences in Zmiz1 impacts on islet mass. While Zmiz1 interacts with the androgen receptor (AR), male mice with specifical deletion of the *Ar* in β-cells showed no obvious defect in islet mass or mass expansion upon metabolic stress (Navarro et al., 2016). The HFD-fed female mice were more resistant to the development of insulin resistance and glucose intolerance, consistent with previous reports that female mice are protected against HFD-induced metabolic changes (Lin et al., 2021; Pettersson et al., 2012), and as such islet mass expanded very little in the female controls.

In summary, our findings demonstrate that ZMIZ1 is crucial for β-cell function and glucose homeostasis and suggest a speculative model by which ZMIZ1 controls a dynamic transcriptional network that governs β-cell maturity and function. Future studies aimed at understanding how ZMIZ1 maintains β-cell functional maturity will be crucial for filling major knowledge gap in current β-cell differentiation protocols for cell replacement therapy.

## Acknowledgements

The University of Alberta is situated on Treaty 6 territory, traditional lands of First Nations and Métis people. We thank Sameena Nawaz (Oxford) for assistance with RNA preparation. We thank the Human Organ Procurement and Exchange (HOPE) program and Trillium Gift of Life Network (TGLN) for their work in procuring human donor pancreas for research, and James Lyon (Edmonton) and Austin Bautista (Edmonton) for their work in human islet isolation at the Alberta Diabetes Institute IsletCore (www.isletcore.ca). We especially thank the organ donors and their families for their kind gift in support of diabetes research.

## Author Contributions

TAA: Performed experiments and analyzed data. Wrote the manuscript.

NAJK: Performed experiments and analyzed data. Wrote and edited the manuscript.

NS, AFS, VR, AJ, HS, MF, JEMF: Performed experiments and analyzed data.

ZS: Designed and generated the floxed Zmiz1 mouse line

ALG: Conceived the study and oversaw the research; Edited the manuscript

PEM: Conceived the study, oversaw the research, edited the manuscript, and acts as guarantor.

## References

1. Ahrén, B., Winzell, M. S., & Pacini, G. (2008). The augmenting effect on insulin secretion by oral versus intravenous glucose is exaggerated by high-fat diet in mice. Journal of Endocrinology, 197(1), 181–187.

2. Ait-Lounis, A., Bonal, C., Seguín-Estévez, Q., Schmid, C. D., Bucher, P., Herrera, P. L., Durand, B., Meda, P., & Reith, W. (2010). The transcription factor Rfx3 regulates beta-cell differentiation, function, and glucokinase expression. Diabetes, 59(7), 1674–1685.

3. Balwierz, P. J., Pachkov, M., Arnold, P., Gruber, A. J., Zavolan, M., & van Nimwegen, E. (2014). ISMARA: Automated modeling of genomic signals as a democracy of regulatory motifs. Genome Research, 24(5), 869–884.

4. Bartolome, A., Zhu, C., Sussel, L., & Pajvani, U. B. (2019). Notch signaling dynamically regulates adult β cell proliferation and maturity. The Journal of Clinical Investigation, 129(1), 268–280.

5. Beliakoff, J., Lee, J., Ueno, H., Aiyer, A., Weissman, I. L., Barsh, G. S., Cardiff, R. D., & Sun, Z. (2008). The PIAS-like protein Zimp10 is essential for embryonic viability and proper vascular development. Molecular and Cellular Biology, 28(1), 282–292.

6. Castillo-Castellanos, F., Ramírez, L., & Lomelí, H. (2021). Zmiz1a zebrafish mutants have defective erythropoiesis, altered expression of autophagy genes, and a deficient response to vitamin D. Life Sciences, 284, 119900.

7. Dobin, A., & Gingeras, T. R. (2015). Mapping RNA-seq reads with STAR. Current Protocols in Bioinformatics, 51(1).

8. Dror, V., Nguyen, V., Walia, P., Kalynyak, T. B., Hill, J. A., & Johnson, J. D. (2007). Notch signalling suppresses apoptosis in adult human and mouse pancreatic islet cells. Diabetologia, 50(12), 2504–2515.

9. Fajas, L., Annicotte, J.-S., Miard, S., Sarruf, D., Watanabe, M., & Auwerx, J. (2004). Impaired pancreatic growth, beta cell mass, and beta cell function in E2F1 (-/-)mice. The Journal of Clinical Investigation, 113(9), 1288–1295.

10. Fidalgo, M., Shekar, P. C., Ang, Y.-S., Fujiwara, Y., Orkin, S. H., & Wang, J. (2011). Zfp281 functions as a transcriptional repressor for pluripotency of mouse embryonic stem cells. Stem Cells (Dayton, Ohio), 29(11), 1705–1716.

11. Gupta, D., Jetton, T. L., LaRock, K., Monga, N., Satish, B., Lausier, J., Peshavaria, M., & Leahy, J. L. (2017). Temporal characterization of β cell-adaptive and -maladaptive mechanisms during chronic high-fat feeding in C57BL/6NTac mice. Journal of Biological Chemistry, 292(30), 12449–12459.

12. Gurzov, E. N., Barthson, J., Marhfour, I., Ortis, F., Naamane, N., Igoillo-Esteve, M., Gysemans, C., Mathieu, C., Kitajima, S., Marchetti, P., Ørntoft, T. F., Bakiri, L., Wagner, E. F., & Eizirik, D. L. (2012). Pancreatic β-cells activate a JunB/ATF3-dependent survival pathway during inflammation. Oncogene, 31(13), 1723–1732.

13. Juliana, C. A., Yang, J., Cannon, C. E., Good, A. L., Haemmerle, M. W., & Stoffers, D. A. (2018). A PDX1-ATF transcriptional complex governs β cell survival during stress. Molecular Metabolism, 17, 39–48.

14. Keller, M. P., Paul, P. K., Rabaglia, M. E., Stapleton, D. S., Schueler, K. L., Broman, A. T., Ye, S. I., Leng, N., Brandon, C. J., Neto, E. C., Plaisier, C. L., Simonett, S. P., Kebede, M. A., Sheynkman, G. M., Klein, M. A., Baliga, N. S., Smith, L. M., Broman, K. W., Yandell, B. S., … Attie, A. D. (2016). The transcription factor Nfatc2 regulates β-cell proliferation and genes associated with type 2 diabetes in mouse and human islets. PLoS Genetics, 12(12), e1006466.

15. Kim-Muller, J. Y., Fan, J., Kim, Y. J. R., Lee, S.-A., Ishida, E., Blaner, W. S., & Accili, D. (2016). Aldehyde dehydrogenase 1a3 defines a subset of failing pancreatic β cells in diabetic mice. Nature Communications, 7, 12631.

16. Kitakaze, K., Oyadomari, M., Zhang, J., Hamada, Y., Takenouchi, Y., Tsuboi, K., Inagaki, M., Tachikawa, M., Fujitani, Y., Okamoto, Y., & Oyadomari, S. (2021). ATF4-mediated transcriptional regulation protects against β-cell loss during endoplasmic reticulum stress in a mouse model. Molecular Metabolism, 54, 101338.

17. Krentz, N. A. J., & Gloyn, A. L. (2020). Insights into pancreatic islet cell dysfunction from type 2 diabetes mellitus genetics. Nature Reviews. Endocrinology, 16(4), 202–212.

18. Ku, H.-C., & Cheng, C.-F. (2020). Master regulator activating transcription factor 3 (ATF3) in metabolic homeostasis and cancer. Frontiers in Endocrinology, 11, 556.

19. Lee, J., Beliakoff, J., & Sun, Z. (2007). The novel PIAS-like protein hZimp10 is a transcriptional co-activator of the p53 tumor suppressor. Nucleic Acids Research, 35(13), 4523–4534.

20. Li, X., Thyssen, G., Beliakoff, J., & Sun, Z. (2006). The novel PIAS-like protein hZimp10 enhances Smad transcriptional activity. The Journal of Biological Chemistry, 281(33), 23748–23756.

21. Liao, Y., Smyth, G. K., & Shi, W. (2014). featureCounts: An efficient general purpose program for assigning sequence reads to genomic features. Bioinformatics, 30(7), 923–930.

22. Lin, H., Smith, N., Spigelman, A. F., Suzuki, K., Ferdaoussi, M., Alghamdi, T. A., Lewandowski, S. L., Jin, Y., Bautista, A., Wang, Y. W., Manning Fox, J. E., Merrins, M. J., Buteau, J., & MacDonald, P. E. (2021). β-cell knockout of SENP1 reduces responses to incretins and worsens oral glucose tolerance in high-fat diet-fed mice. Diabetes, 70(11), 2626–2638.

23. Love, M. I., Huber, W., & Anders, S. (2014). Moderated estimation of fold change and dispersion for RNA-seq data with DESeq2. Genome Biology, 15(12), 550.

24. Lyon, J., F Spigelman, A., E Macdonald, P., & E Manning Fox, J. (2019). ADI IsletCore protocols for the isolation, assessment and cryopreservation of human pancreatic islets of Langerhans for research purposes v1. protocols.io. doi.org/10.17504/protocols.io.x3mfqk6

25. Mahajan, A., Spracklen, C. N., Zhang, W., Ng, M. C. Y., Petty, L. E., Kitajima, H., Yu, G. Z., Rüeger, S., Speidel, L., Kim, Y. J., Horikoshi, M., Mercader, J. M., Taliun, D., Moon, S., Kwak, S.-H., Robertson, N. R., Rayner, N. W., Loh, M., Kim, B.-J., … Morris, A. P. (2022). Multi-ancestry genetic study of type 2 diabetes highlights the power of diverse populations for discovery and translation. Nature Genetics, 54(5), 560–572.

26. Mahajan, A., Taliun, D., Thurner, M., Robertson, N. R., Torres, J. M., Rayner, N. W., Payne, A. J., Steinthorsdottir, V., Scott, R. A., Grarup, N., Cook, J. P., Schmidt, E. M., Wuttke, M., Sarnowski, C., Mägi, R., Nano, J., Gieger, C., Trompet, S., Lecoeur, C., … McCarthy, M. I. (2018). Fine-mapping type 2 diabetes loci to single-variant resolution using high-density imputation and islet-specific epigenome maps. Nature Genetics, 50(11), 1505–1513.

27. Mattis, K. K., & Gloyn, A. L. (2020). From genetic association to molecular mechanisms for islet-cell dysfunction in type 2 diabetes. Journal of Molecular Biology, 432(5), 1551–1578.

28. Navarro, G., Xu, W., Jacobson, D. A., Wicksteed, B., Allard, C., Zhang, G., De Gendt, K., Kim, S. H., Wu, H., Zhang, H., Verhoeven, G., Katzenellenbogen, J. A., & Mauvais-Jarvis, F. (2016). Extranuclear actions of the androgen receptor enhance glucose-stimulated insulin secretion in the male. Cell Metabolism, 23(5), 837–851.

29. Pettersson, U. S., Waldén, T. B., Carlsson, P.-O., Jansson, L., & Phillipson, M. (2012). Female mice are protected against high-fat diet induced metabolic syndrome and increase the regulatory T cell population in adipose tissue. PloS One, 7(9), e46057.

30. Pieraccioli, M., Nicolai, S., Pitolli, C., Agostini, M., Antonov, A., Malewicz, M., Knight, R. A., Raschellà, G., & Melino, G. (2018). ZNF281 inhibits neuronal differentiation and is a prognostic marker for neuroblastoma. Proceedings of the National Academy of Sciences of the United States of America, 115(28), 7356–7361.

31. Pinnell, N., Yan, R., Cho, H. J., Keeley, T., Murai, M. J., Liu, Y., Alarcon, A. S., Qin, J., Wang, Q., Kuick, R., Elenitoba-Johnson, K. S. J., Maillard, I., Samuelson, L. C., Cierpicki, T., & Chiang, M. Y. (2015). The PIAS-like coactivator Zmiz1 Is a direct and selective cofactor of Notch1 in T cell development and leukemia. Immunity, 43(5), 870–883.

32. Rady, B., Chen, Y., Vaca, P., Wang, Q., Wang, Y., Salmon, P., & Oberholzer, J. (2013). Overexpression of E2F3 promotes proliferation of functional human β cells without induction of apoptosis. Cell Cycle (Georgetown, Tex.), 12(16), 2691–2702.

33. Ritchie, M. E., Phipson, B., Wu, D., Hu, Y., Law, C. W., Shi, W., & Smyth, G. K. (2015). Limma powers differential expression analyses for RNA-sequencing and microarray studies. Nucleic Acids Research, 43(7), e47–e47.

34. Salinno, C., Büttner, M., Cota, P., Tritschler, S., Tarquis-Medina, M., Bastidas-Ponce, A., Scheibner, K., Burtscher, I., Böttcher, A., Theis, F. J., Bakhti, M., & Lickert, H. (2021). CD81 marks immature and dedifferentiated pancreatic β-cells. Molecular Metabolism, 49, 101188.

35. Salinno, C., Cota, P., Bastidas-Ponce, A., Tarquis-Medina, M., Lickert, H., & Bakhti, M. (2019). β-cell maturation and identity in health and disease. International Journal of Molecular Sciences, 20(21), E5417.

36. Sharma, M., Li, X., Wang, Y., Zarnegar, M., Huang, C.-Y., Palvimo, J. J., Lim, B., & Sun, Z. (2003). HZimp10 is an androgen receptor co-activator and forms a complex with SUMO-1 at replication foci. The EMBO Journal, 22(22), 6101–6114.

37. Shuai, K., & Liu, B. (2005). Regulation of gene-activation pathways by PIAS proteins in the immune system. Nature Reviews. Immunology, 5(8), 593–605.

38. Simonett, S. P., Shin, S., Herring, J. A., Bacher, R., Smith, L. A., Dong, C., Rabaglia, M. E., Stapleton, D. S., Schueler, K. L., Choi, J., Bernstein, M. N., Turkewitz, D. R., Perez-Cervantes, C., Spaeth, J., Stein, R., Tessem, J. S., Kendziorski, C., Keleş, S., Moskowitz, I. P., … Attie, A. D. (2021). Identification of direct transcriptional targets of NFATC2 that promote β cell proliferation. The Journal of Clinical Investigation, 131(21), e144833.

39. Smith, N., F Spigelman, A., Lin, H., & E Macdonald, P. (2018). Mouse pancreatic islet isolation v1. protocols.io. doi.org/10.17504/protocols.io.sqaedse

40. Smith, N., Ferdaoussi, M., Lin, H., & E Macdonald, P. (2018). Oral glucose tolerance test in mouse v1. protocols.io. doi.org/10.17504/protocols.io.ujjeukn

41. Smith, N., Ferdaoussi, M., Lin, H., & E Macdonald, P. (2019a). Insulin tolerance test in mouse v1. protocols.io. doi.org/10.17504/protocols.io.wxjffkn

42. Smith, N., Ferdaoussi, M., Lin, H., & E Macdonald, P. (2019b). IP glucose tolerance test in mouse v1. protocols.io. doi.org/10.17504/protocols.io.wxhffj6

43. Smith, N., Lin, H., Ferdaoussi, M., & E Macdonald, P. (2018). Purification of mouse pancreatic islets using histopaque gradient centrifugation v1. protocols.io. doi.org/10.17504/protocols.io.u7ueznw

44. Soriano, P. (1999). Generalized lacZ expression with the ROSA26 Cre reporter strain. Nature Genetics, 21(1), 70–71.

45. Spracklen, C. N., Horikoshi, M., Kim, Y. J., Lin, K., Bragg, F., Moon, S., Suzuki, K., Tam, C. H. T., Tabara, Y., Kwak, S.-H., Takeuchi, F., Long, J., Lim, V. J. Y., Chai, J.-F., Chen, C.-H., Nakatochi, M., Yao, J., Choi, H. S., Iyengar, A. K., … Sim, X. (2020). Identification of type 2 diabetes loci in 433,540 East Asian individuals. Nature, 582(7811), 240–245.

46. Thomsen, S. K., Ceroni, A., van de Bunt, M., Burrows, C., Barrett, A., Scharfmann, R., Ebner, D., McCarthy, M. I., & Gloyn, A. L. (2016). Systematic functional characterization of candidate causal genes for type 2 diabetes risk variants. Diabetes, 65(12), 3805–3811.

47. Thorens, B., Tarussio, D., Maestro, M. A., Rovira, M., Heikkilä, E., & Ferrer, J. (2015). Ins1(Cre) knock-in mice for beta cell-specific gene recombination. Diabetologia, 58(3), 558–565.

48. Thurner, M., van de Bunt, M., Torres, J. M., Mahajan, A., Nylander, V., Bennett, A. J., Gaulton, K. J., Barrett, A., Burrows, C., Bell, C. G., Lowe, R., Beck, S., Rakyan, V. K., Gloyn, A. L., & McCarthy, M. I. (2018). Integration of human pancreatic islet genomic data refines regulatory mechanisms at Type 2 Diabetes susceptibility loci. ELife, 7, e31977.

49. van de Bunt, M., Manning Fox, J. E., Dai, X., Barrett, A., Grey, C., Li, L., Bennett, A. J., Johnson, P. R., Rajotte, R. V., Gaulton, K. J., Dermitzakis, E. T., MacDonald, P. E., McCarthy, M. I., & Gloyn, A. L. (2015). Transcript expression data from human islets links regulatory signals from genome-wide association studies for type 2 diabetes and glycemic traits to their downstream effectors. PLoS Genetics, 11(12), e1005694.

50. van de Geijn, B., McVicker, G., Gilad, Y., & Pritchard, J. K. (2015). WASP: Allele-specific software for robust molecular quantitative trait locus discovery. Nature Methods, 12(11), 1061–1063.

51. Viñuela, A., Varshney, A., van de Bunt, M., Prasad, R. B., Asplund, O., Bennett, A., Boehnke, M., Brown, A. A., Erdos, M. R., Fadista, J., Hansson, O., Hatem, G., Howald, C., Iyengar, A. K., Johnson, P., Krus, U., MacDonald, P. E., Mahajan, A., Manning Fox, J. E., … McCarthy, M. I. (2020). Genetic variant effects on gene expression in human pancreatic islets and their implications for T2D. Nature Communications, 11(1), 4912.

52. Wortel, I. M. N., van der Meer, L. T., Kilberg, M. S., & van Leeuwen, F. N. (2017). Surviving stress: Modulation of ATF4-mediated stress responses in normal and malignant cells. Trends in Endocrinology and Metabolism: TEM, 28(11), 794–806.

53. Yamane, S., Harada, N., & Inagaki, N. (2016). Mechanisms of fat-induced gastric inhibitory polypeptide/glucose-dependent insulinotropic polypeptide secretion from K cells. Journal of Diabetes Investigation, 7(S1), 20–26.

54. Zmuda, E. J., Qi, L., Zhu, M. X., Mirmira, R. G., Montminy, M. R., & Hai, T. (2010). The roles of ATF3, an adaptive-response gene, in high-fat-diet-induced diabetes and pancreatic beta-cell dysfunction. Molecular Endocrinology (Baltimore, Md.), 24(7), 1423–1433.

