## Supplemental Figures for "Zmiz1 is required for mature β-cell function and mass expansion upon high fat feeding"

**Supplemental Figure Legends**

**Suppl Figure 1. Distribution of donor characteristics in carriers of the T2D-associated rs703972 and rs12571751 risk alleles in *ZMIZ1.*** Sex **(A)**, age **(B)**, body mass index **(C)**, and glycated hemoglobin (HbA1c) **(D)** in donors of different genotype at SNPs rs703972 and rs12571751. Non-risk allele donors possess the rs703972 C allele (CC, n=37) and the rs12571751 G allele (GG, n=39) whereas heterozygous (rs703972 CG, n=126; rs12571751 GA, n=127) and homozygous (rs703972 GG, n=69; rs12571751 AA, n=66) donors are carriers of the T2D risk alleles. **(E-F)** Assessment of the impact of the lead SNP (rs1317617) at the second *ZMIZ1* signal on total insulin content (**E**) and insulin secretion in response to low glucose (1 mM) or high glucose (16.7 mM) (**F**) of donors possessing at rs1317617 the A allele (AA, n=18), heterozygous (GA, n=73) or carriers of the G allele (GG, n=124).

**Suppl Figure 2. Cre activity and insulin tolerance in β-cell specific Zmiz1 knockout mice. (A)** Enzymatic X-gal staining of mouse pancreatic cryosections from Ins1-Cre$^{-}$R26R$^{fl/+}$ or Ins1-Cre$^{+}$R26R$^{fl/+}$. **(B-C)** Insulin tolerance in female (B) and male (C) control (Zmiz1$^{Ctrl}$), heterozygous (Zmiz1$^{\beta Het}$) and β-cell-specific knockout (Zmiz1$^{\beta KO}$) mice. n=6-8 per group.

**Suppl Figure 3. Additional data on Zmiz1$^{\beta KO}$ mice after high fat diet (HFD). (A)** IP glucose tolerance and **(B)** insulin tolerance of female control (Zmiz1$^{Ctrl}$), heterozygous (Zmiz1$^{\beta Het}$) and β-cell-specific knockout (Zmiz1$^{\beta KO}$) mice after an 8-week HFD (n=6-9 mice per group). **(C)** Fasting plasma insulin in female (Zmiz1$^{Ctrl}$, Zmiz1$^{\beta Het}$ and Zmiz1$^{\beta KO}$ mice after an 8-week HFD (n=5-10 mice per group). **(D-E)** Representative insulin immunostaining (D) and quantification of β-cell mass (E) in pancreas of female Zmiz1$^{Ctrl}$, Zmiz1$^{\beta Het}$ and Zmiz1$^{\beta KO}$ mice after an 8-week HFD (n=4-7 mice per group) (scale bars= 100 μm). **(F-G)** Glucose-stimulated insulin secretion from islets of female (F) and male (G) Zmiz1$^{Ctrl}$ and Zmiz1$^{\beta KO}$ mice after an 8-week HFD (n=5-9 mice per group). Data are mean ± SEM. $^{*}P<0.05$ and $^{**}P<0.01$ by one-way ANOVA followed by Tukey's multiple comparisons test.

#### Suppl Figure 1

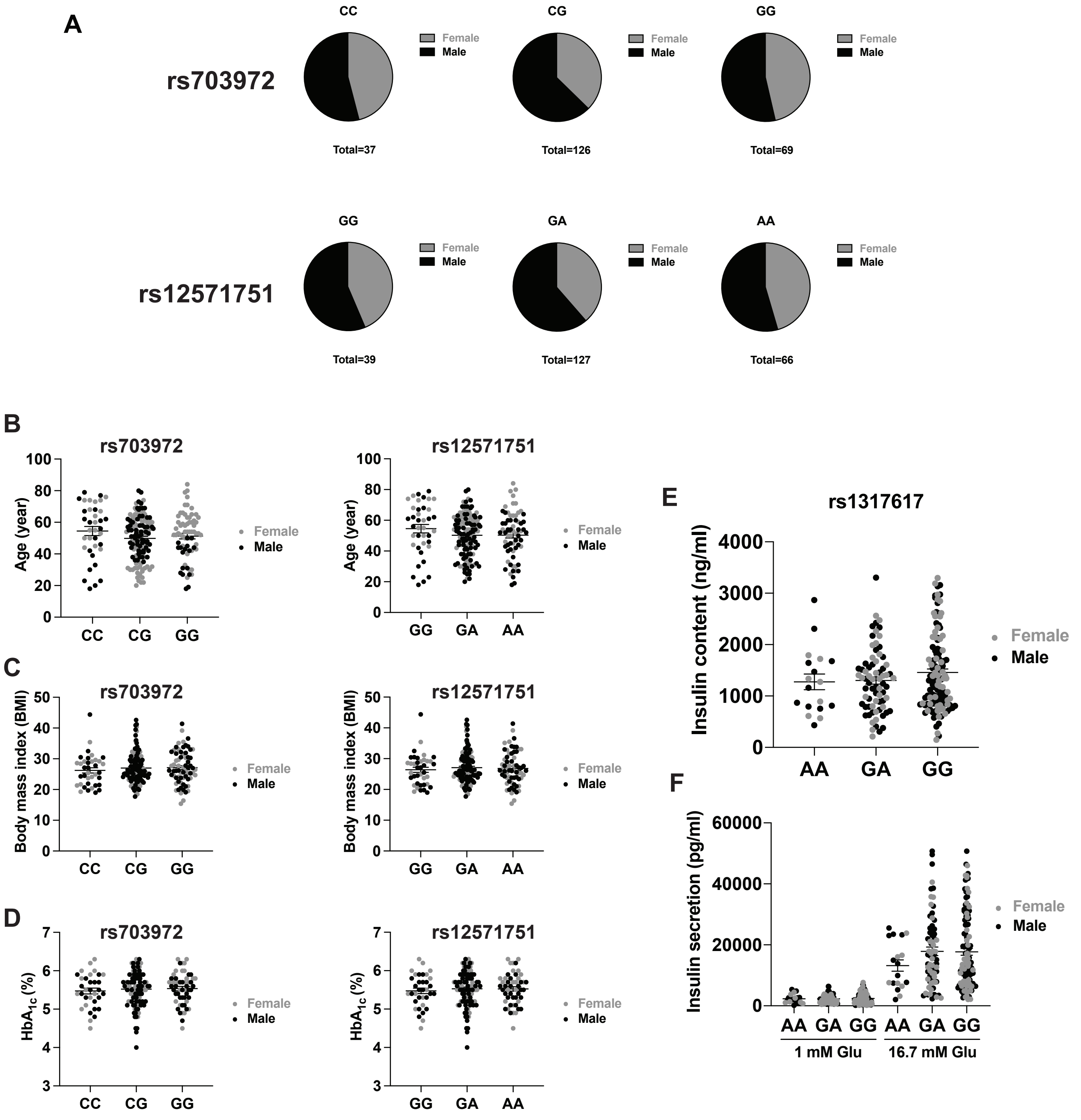

### Suppl Figure 2

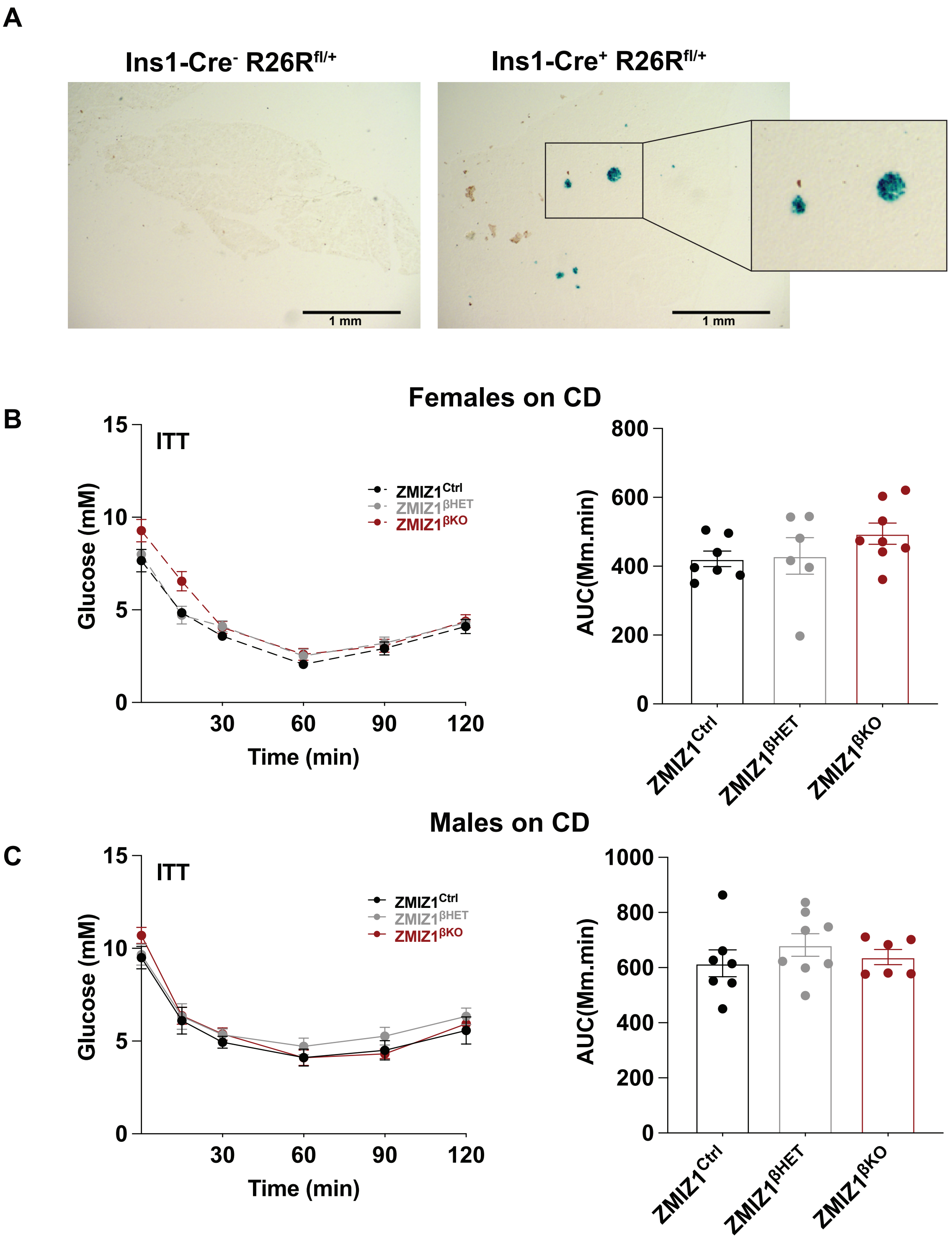

### Suppl Figure 3

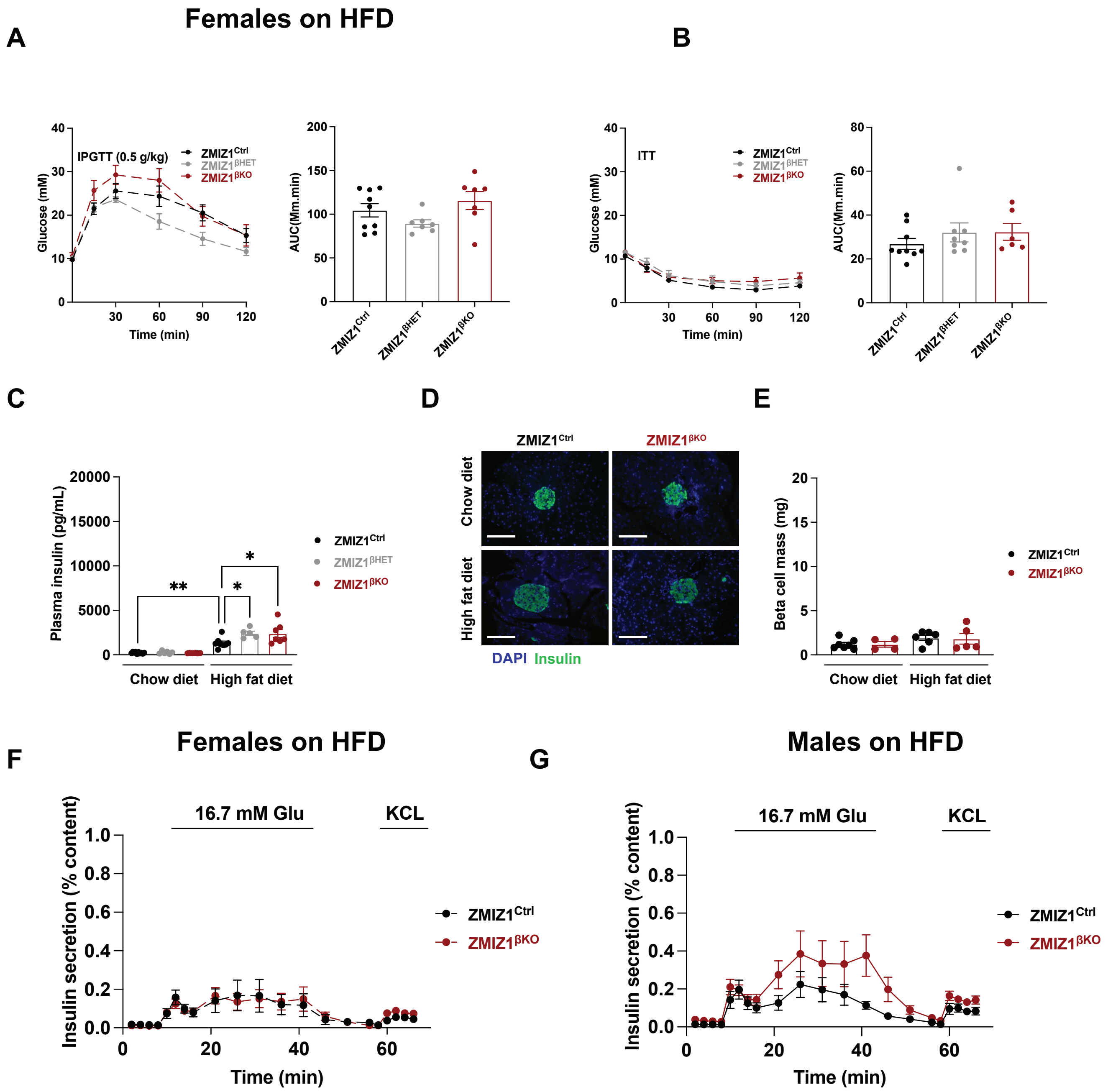
